# WorMachine: Machine Learning-Based Phenotypic Analysis Tool for Worms

**DOI:** 10.1101/211359

**Authors:** Adam Hakim, Yael Mor, Itai Antoine Toker, Amir Levine, Yishai Markovitz, Oded Rechavi

## Abstract

While *Caenorhabditis elegans* nematodes are powerful model organisms, quantification of visible phenotypes is still often labor-intensive, biased, and error-prone. We developed “WorMachine”, a three-step MATLAB-based image analysis software that allows automated identification of *C. elegans* worms, extraction of morphological features, and quantification of fluorescent signals. The program offers machine learning techniques which should aid in studying a large variety of research questions. We demonstrate the power of WorMachine using five separate assays: scoring binary and continuous sexual phenotypes, quantifying the effects of different RNAi treatments, and measuring intercellular protein aggregation. Thus, WorMachine is a “quick and easy”, high-throughput, automated, and unbiased analysis tool for measuring phenotypes.

## Background

*Caenorhabditis elegans* nematodes are powerful genetic model organisms which are instrumental for research of a wide-range of biological questions. It is relatively simple to grow *C. elegans* under tightly regulated environmental conditions, and since the worm has a short generation time (3 days), and a large brood size (±250), large sample sizes and statistical power are easily obtained. In many cases, however, when phenotypic features are examined, the advantage of having a large sample size comes with great cost, because of the need to manually analyze the features of interest in the tested animals. While in the last several years very useful programs for quantifying *C. elegans*’ phenotypes from still images were released, such as WormSizer [1], Fiji [2], and QuantWorm [3], the analysis process is not fully automated, and not all informative features can be analyzed simultaneously in one package.

We created WorMachine as a fast, friendly, and high-throughput image processing platform. WorMachine enables automated calculation of many morphological and fluorescence features, and accessible machine learning techniques for higher-level features-based analysis, such as classification and phenotype scoring. WorMachine is entirely MATLAB-based, and combines the capabilities of different programs into one software; the user-friendly interface was designed to suit investigators with no background in MATLAB, image processing, or machine learning, and requires no additional plugins or installations. WorMachine is not limited to any specific image format, resolution, acquisition software, or microscope.

## Implementation

WorMachine’s workflow includes three sequential programs: Image Processor, Feature Extractor, and Machine Learner **(Fig. 1)**. WorMachine’s codes, demonstration video, and a sample TIFF image to try the program with, are accessible through the supplied manual, available in the **Supplementary Materials**.

**Figure 1.**
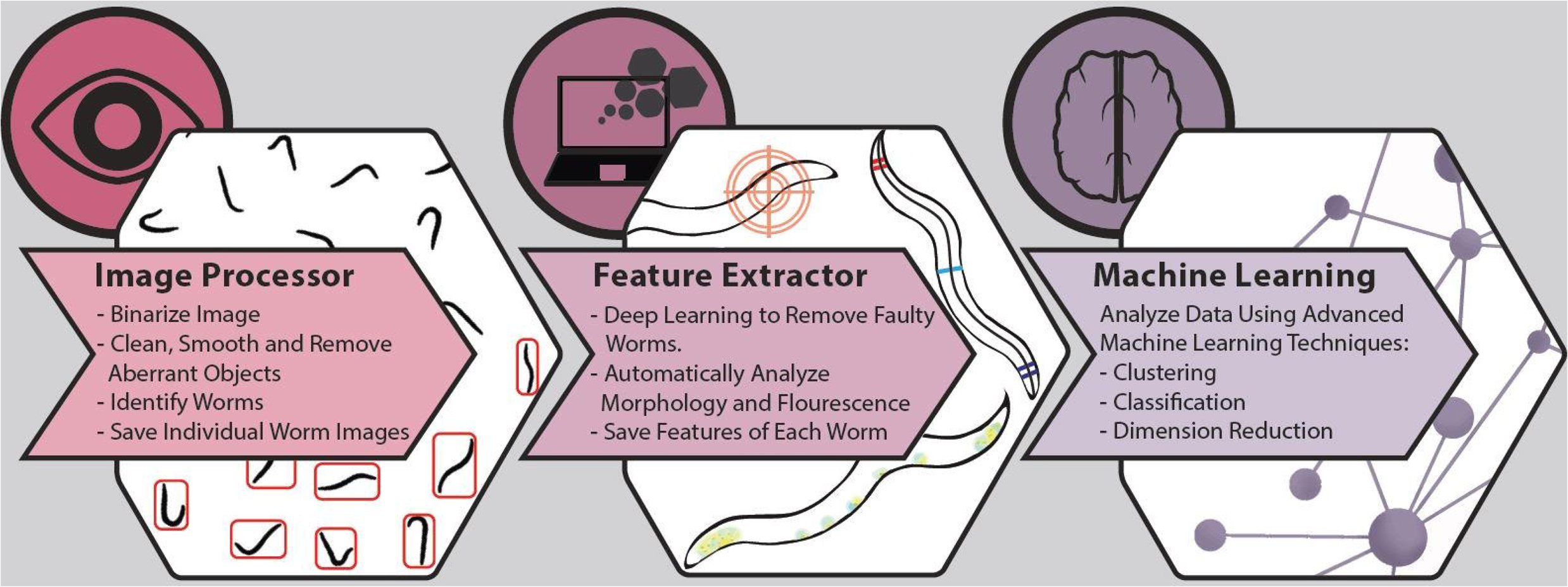
WorMachine workflow. The software includes three sequential programs: An Image Processor, a Feature Extractor, and a Machine Learner.

The *Image Processor* uses still bright-field images of worm plates as input (acquired using any typical image acquisition microscope). Fluorescent images can similarly be analyzed together with overlapping bright-field acquisitions. The image acquisition procedures and parameters which enabled optimal processing of images in our hands are detailed in the ‘Online Methods’ section. Identifying “real” worms from other elements is normally a painstaking stage that delays image analysis. The Image processor of WorMachine binarizes, identifies, and crops individual worms from the original image automatically.

The cropped worm masks are then loaded to the *Feature Extractor*, where the worms’ morphological and fluorescent features are analyzed individually. During this stage of the analysis, potentially faulty and damaged worm images are flagged by a deep learning network designed particularly for this task, and made available for the investigator’ review (in case these should be removed). The Feature Extractor also enables tagging different worms with labels according to the user’s needs, such as assigning worms to different conditions or groups. For example, the worms’ sex can be labeled, and this information can be used for creating a training dataset for later classification.

Finally, the *Machine Learner* builds on the obtained features and labels to conduct binary classification using Support Vector Machine, or visualization and scoring of high-dimensional data and dimensionality reduction using “Principal Component Analysis” (PCA) or "t-Distributed Stochastic Neighbor Embedding" (t-SNE) [4,5].

## Image Processor

WorMachine is best suited to handle TIFF images with one or multiple channels, which can have a maximum size of about 500 megabytes, depending on the memory of the user’s computer. However, it also supports a wide range of additional formats used by biologists, as it incorporates the <u>Bio-Formats Library</u> [6] for image reading. Imported images are automatically gray-scaled and contrast-adjusted to accentuate the differences between worms and the background, and accordingly to improve the detection of nematodes. The program generates a binary mask based on the imported image, using adaptive local thresholding [7], which is then “cleaned” and segmented using MATLAB’s own image processing toolbox (for details, see “Online Methods”). Individual worms within a likely size are automatically identified, smoothed, filled, and cleaned, using standard image processing procedures included in MATLAB’s Image processing toolbox. All individual worm images are numbered and saved to a folder based on their respective image channel (BF, Fluorescence, Masks, Etc.). This procedure may be applied automatically on multiple images, using the “Batch Analysis” option (see Manual and demo movie, in the Supplementary Materials).

## Feature Extractor

Once individual worm masks are imported into this program, the analysis may be performed on all channels in parallel. All morphological and fluorescence measurements currently available in WorMachine are detailed in **Table 1,** and each extraction technique is detailed in the “Online Methods” section. After extraction, objects which deviate in area size, length, or skeleton disfigurement are flagged for manual inspection, together with images identified as “noise” by the deep-learning network. Thus, the user may further clean and refine her database. The proportion of worms excluded by this step is detailed in **Supplementary Table 1**.

**Table 1.** Morphological and fluorescent features and their calculation methods.

| **Feature** | **Calculation** |
| --- | --- |
| ***Area*** | *Number of pixels in the worm's area X pixel height*<br>*X pixel width* |
| ***Length*** | *Number of pixels in the worm's skeleton X pixel height* |
| ***Thickness*** | *Area/Length* |
| ***Midwidth*** | *Number of pixels in the worm's center width X pixel height* |
| ***Head & Tail Diameter Ratios*** | $\frac{\text{Diameter at thickest section 10\% from edge}}{\text{Diameter at slimmest section 10\% – 20\% from edge}}$ |
| ***Head BF*** | *Mean Brightness of a polygon within the worm's head* |
| ***Tail BF*** | *Mean Brightness of a polygon within the worm's tail* |
| ***Peaks Count*** | *2d Local Maximum of Fluorescent Intensities* |
| ***Mean & STD of Peaks*** | *Average & Standard Deviation of peak intensities* |
| ***CTCF*** | *Mean Worm Fluorescence X Worm Area*<br>*– Mean Background Fluorescence X Worm Area* |
| ***RID*** | *Sum of Worm Fluorescence in its Area*<br>*– Mean Background Fluorescence X Worm Area* |

## Machine Learner

At this stage, data extracted from the previous stages can be analyzed with different Machine-Learning techniques. First, users may visually review and select features relevant to their analysis. Next, the user can choose between two techniques: (1) SVM for binary classification based on supplied or user-generated training data, (2) High-dimensionality visualization and scoring of complex phenotypes based on various features using PCA or t-SNE. The algorithms and their use are detailed in the “Online Methods” section, and examples for different applications are provided in the “Results” section.

## Results

We examined WorMachine’s ability to facilitate analyses of multiple different phenotypes. First, we describe the use of the software for classification of worm populations based on binary sex phenotypes (males or hermaphrodites). Then we demonstrate, using a mutant that displays a continuous sexual phenotype, that WorMachine can accurately create a common scale of worm masculinization. Next, we show how the software can be used to quantify RNAi-induced gene silencing, protein aggregation, and puncta distribution.

## Binary classification of the worms’ sex

*C.elegans* nematodes have two sexes – the great majority of the worms are self-fertilizing XX hermaphrodites, and a small minority (0.1%- 0.2%) are X0 males [8]. WorMachine can be used to calculate in a high-throughput and precise manner the sex ratios in different strains and conditions. To distinguish between the sexes WorMachine uses morphological and brightness features that differentiate between hermaphrodites and males, and also, when fluorescent reporters are available, sex-specific expression patterns. The mutant worms that we used here (*him-5(e1467); zdIs13 [tph-1p::GFP] IV])*, segregate many males, and express GFP in the serotonergic neurons. Mutations in *him-5* increase the frequency of XO males (to about 30%) by elevating the frequency of X chromosome nondisjunction [9]. The *tph1p::GFP* transgene allows distinguishing the worms’ sex as it drives GFP expression in males-specific and hermaphrodites-specific neurons: the hermaphrodite-specific neuron (HSN), the males ventral cord motor neurons (CPs), and some tale-specific neurons [10,11]. We classified worms based on morphological, brightness, and fluorescent features (**Supplementary Fig. 1**), and reached 98% classification accuracy when we trained on all features using 1800 worms. **Figure 2** displays the true positive rates of the machine-learning program, based on training sets of different sizes (30 to 2000 worms), with or without taking advantage of the sex-specific fluorescence pattern.

**Figure 2.**
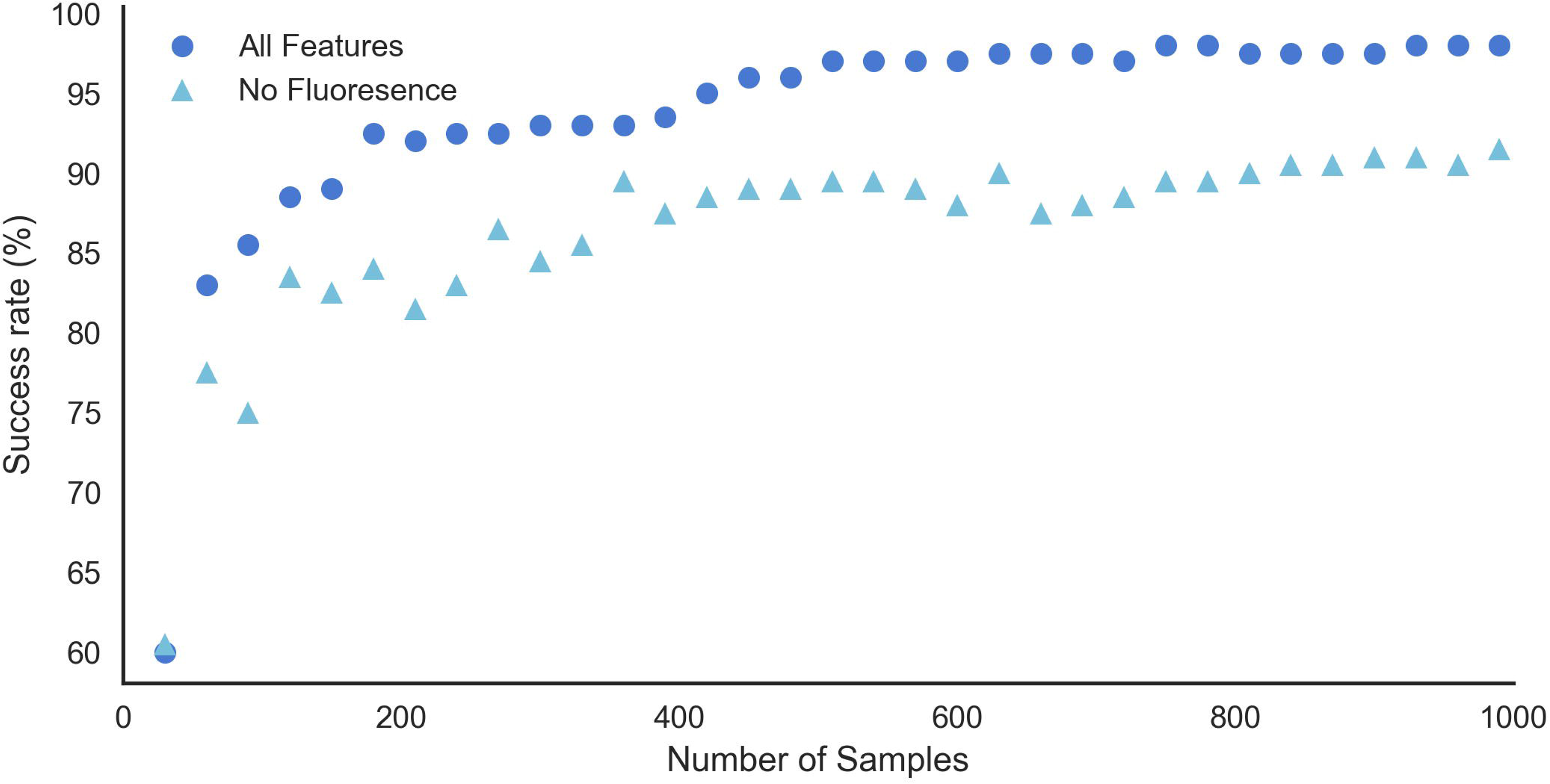
Success rates in classifying the worms’ sex as a function of the number of worms used for training. Shown are results when the fluorescence of a reporter expressed in sex-specific neurons was taken into account (dark blue, dots), and when only morphological and brightness features were considered (light blue, triangles). The classification is based on these morphological features: Head BF, Tail BF, Area, Length, Midwidth, Thickness, Tail Ratio, and on these fluorescent features: Peak number (count), CTCF, and Mean peak intensity.

## Quantifying a continuous sex phenotype

Continuous morphological phenotypes are common, and due to their complex nature, their quantification is often challenging. We used the CB5362 strain, which is mutated in the sex determination genes *xol-1* and *tra-2*. These worms display an intersex phenotype, which depends on temperature [12]. We used WorMachine to determine the sexual phenotype (= degree of masculinization) of each worm, based on multiple features: The shape of the tale (angle evaluation) [13], the presence or absence of eggs in the gonad (eggs bearing worms have larger mid-width) the worm's length and area (males are smaller than hermaphrodites), the head and tail brightness (males have darker tales in bright field) (**Supplementary Fig. 1**). We grew CB5362 worms at three different temperatures (15, 20 & 25 degrees Celsius) and imaged them at the first day of adulthood. The program determined the sex of each worm, based on a sexual phenotype spectrum ranging from male to hermaphrodite, using dimensions reduction. The technique yielded scores that were concurrent with previous literature, showing higher masculinity scores for higher temperatures (**Fig 3, Supplementary figures 2,3**).

**Figure 3.**
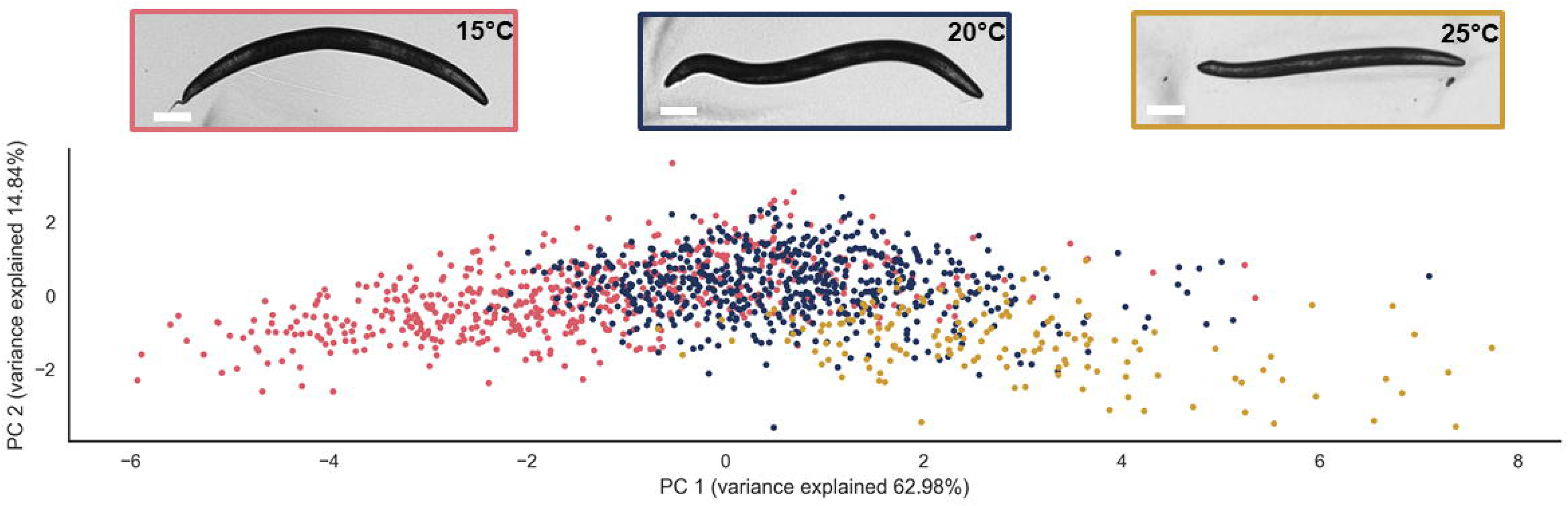
A PCA visualizing the effect of temperature on the sexual phenotype. CB5362 worms were grown in different temperatures and imaged during the first day of adulthood. The PCA was calculated on scaled data and was based on the statistically significance features that distinguish between the sexes (e.g. Area, Length, Midwidth, Thickness, Tail ratio, Head BF, and tail BF, as Supplementary Fig 1.A). Red = 15 degrees Celsius; blue = 20 degrees Celsius; yellow = 25 degrees Celsius. Each dot represents an individual worm. Upper panel, representative worm at each temperature.

## Quantifying RNAi-induced phenotypes

We used WorMachine to quantify the RNAi response of worms fed with anti-*dpy-11* or anti-*mCherry* dsRNA-expressing bacteria.

### anti-dpy-11 RNAi

Knockdown of *dpy-11* results in a “Dumpy” phenotype (lower length) [14]. In addition to using wild-type worms (N2), we examined the RNAi response also in *rrf-3(pk1426)* mutants, which are hypersensitive to RNAi (exhibit an Enhanced RNAi, or Eri, phenotype) (Simmer *et al.*, 2002). As can be seen in **Fig. 4.A**, WorMachine successfully captures the stronger response to RNAi of *rrf-3* mutants, in comparison to N2 wild-types (p<10^−4^).

### Anti-mCherry RNAi

We used worms that express mCherry ubiquitously (EG7841 *oxTi302 [eft-3p::mCherry::tbb-2 3'UTR + Cbr-unc- 119(+)],* [15]. Worms were treated with dsRNA-expressing bacteria grown to different O.D. values (to obtain a gradient of silencing efficiencies), and whole-worm CTCF was measured. As expected, worms exposed to bacteria grown to higher O.D values, showed lower CTCF values which reflect greater levels of silencing (p<10^−4^) (**Fig. 4.B**). The differences in CTCF values (silencing levels) that were measured by WorMachine were comparable to the differences measured when using ImageJ (**Supplementary Fig.4**).

**Figure 4.**
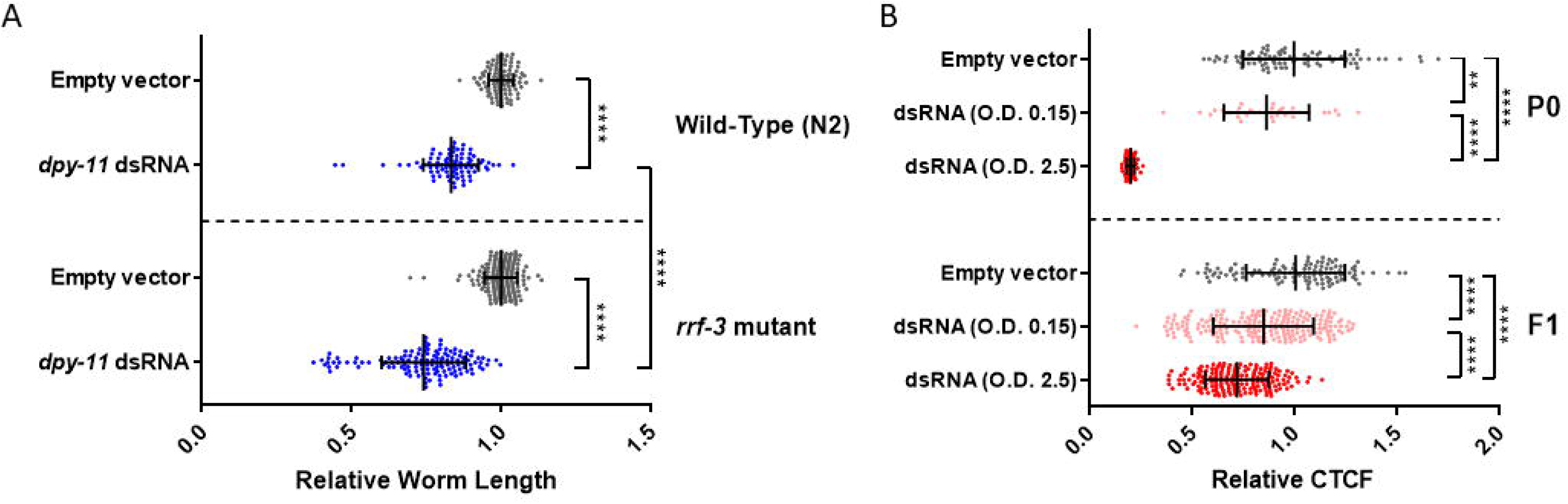
Quantification of RNAi-induced phenotypes. The worms were imaged during the first day of adulthood. The values for each worm were divided by the mean value of the corresponding control group. Each dot represents an individual worm. Bars represent mean ± standard deviation. **** p<10^−4^, ** p<10^−2^. **A.** Quantification of the Dumpy phenotype following *dpy-11* RNAi treatment. N2 worms (upper panel) or *rrf-3* mutants (lower panel) were fed with bacteria expressing an empty vector control or dsRNA complementary to *dpy-11*. **B.** Quantification of fluorescence intensity in whole animals. EG7841 worms expressing mCherry in all somatic cells were fed with bacteria expressing either an empty vector control or dsRNA complementary to mCherry. The RNAi-producing bacteria were grown to the optical density (O.D.) which is indicated. P0 condition: The eggs were laid on RNAi-producing bacteria lawns. F1 condition: The progeny of the RNAi treated worms, that were laid and grown on standard OP50 bacteria.

## Quantification of intercellular protein aggregation

Protein aggregation can be toxic, and is a hallmark of many diseases [16]. The Cohen lab (The Hebrew University) studies the cellular mechanisms of polyglutamine toxicity, and agreed to test whether WorMachine can be useful for quantifying the aggregation of polyglutamine proteins. Importantly, the analysis of this phenotype and the acquisition of the data was done outside of the Rechavi lab, using a different microscope, and by non-Rechavi lab members (Amir Levine, from the Cohen lab). The transgenic AM140 worm strain expresses a polyglutamine protein (35 repeats) tagged with the Yellow Fluorescent Protein (polyQ-YFP) in body wall muscles [17]. These animals form visible polyglutamine puncta that accumulate in an age-dependent manner. The sizes and quantity of these puncta serves as a measure for toxic polyglutamine aggregation [17,18]. The large variability of puncta quantities among worms in a population, and the large differences in puncta sizes within each individual worm, normally requires the collection of large data sets to achieve reproducible and consistent results. WorMachine was able to measure the number and size distributions of polyQ-YFP in a high-throughput manner. The abundance of polyQ35-YFP puncta increases with age (in accordance with the literature), while the relative sizes of polyQ35-YFP puncta decreases **(Fig.5)**. The differences in the number of puncta between the different experimental conditions (different days) that were identified manually were also identified by WorMachine (**Supplementary Fig 4**).

**Figure 5.**
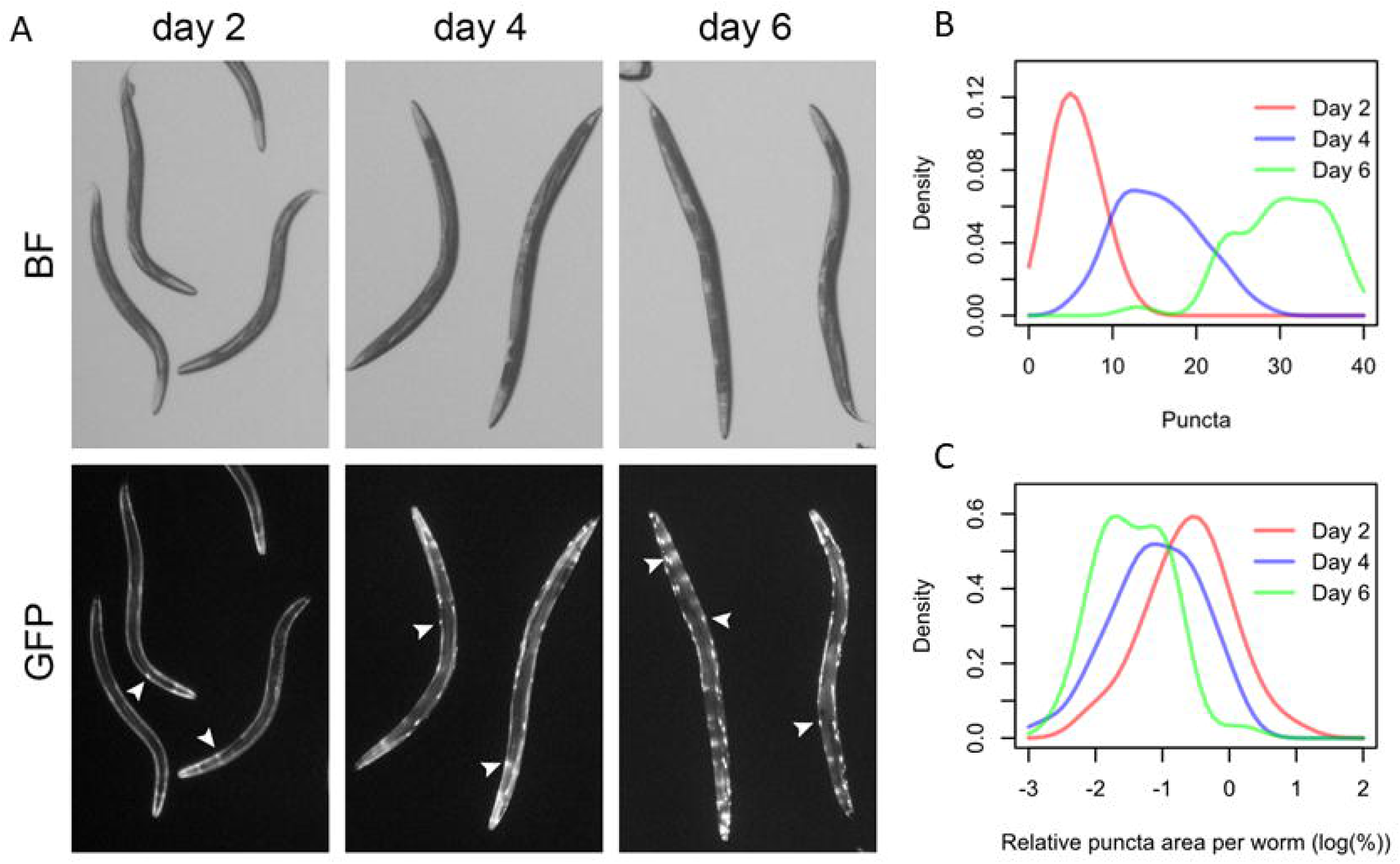
Quantifying intercellular protein aggregation. **A** Representative images of polyQ35-YFP expressing animals at day 2, 4, and 6 of adulthood. BF = bright field illumination. Fluorescent protein aggregations are marked with white arrows. **B.** The abundance of polyQ35-YFP puncta at days 2 (red, n=64), 4 (blue, n=37), and 6 (green, n=42) of adulthood. p<10^−4^. **C.** The relative sizes of polyQ35-YFP puncta per worm on days 2 (red), 4 (blue), and 6 (green) of adulthood. Distributions of relative puncta sizes per worm at all ages, shown as a density plot.

## Conclusions

WorMachine offers the nematode research community an easy-to-used, automated, accurate, and unbiased methodology to analyze morphological and fluorescent worm features from images obtained using standard microscopes. The program analyzes images in three steps, detection of worms, extraction of features, and machine-learning applications. The software has a modular design in each step that is adaptable to user-specific requirements.

Importantly, WorMachine also includes a deep-learning neural network that recognizes, flags, and omits non-suitable objects, to ensure a high level of quality for the users’ generated data. The use of the machine-learning algorithms can uncover new information about the data, that would be hard to reveal using standard analyses. The software is user-friendly, free, and can be edited by any user, as it is implemented in MATLAB. There are many more possible applications for WorMachine, in addition to quantification of the biological features that we analyzed in this paper, and hopefully many users would find this software helpful in the near future.

## Online Methods

### Preparation of Worms for Imaging

Worms were synchronized at each generation using one of two standard ways: (1) Mothers were allowed to lay eggs for a limited span of 24 hours, or (2) by bleaching (“egg-prep”) [19]. Adult worms were washed 3 times to get rid of bacteria (OP50) residues. Worms were left in ~100 microliter of M9 buffer and paralyzed via the addition of Sodium Azide (final concentration of 25-50mM). The paralyzed worms were transferred to imaging plates and then physically separated from each other using a platinum-wire pick. Separation of the worms is important for allowing the software to correctly process each worm individually. The imaging plates were 60mm petri dishes filled with 8mL of modified transparent NGM (2% Agarose, 0.3% NaCl, 5mM K_2_PO_4_, 1mM CaCl_2_, 1mM MgSO_4_).

### Microscopy Image Acquisition

Images were taken on an Olympus IX83 microscope, using fluorescence excitation with a LED light source on two channels: Green Fluorescent Protein (GFP) (FITC/Cy2) and mCherry Fluorescence. For bright field (BF) imaging, a relatively long exposure time was used for trails erasure and contrast was increased to better differentiate between worms and background. Pictures were taken with 4X/0.75 Universal Plan Super Apochromat objective.

For the Protein aggregation assays images were acquired using a Nikon SMZ18 stereoscope fitted with 1X objective, set-up to capture both bright-field illumination and YFP fluorescence.

### *C. elegans* Strains

The *C. elegans* strains employed in this work are as follows: wild-type Bristol N2 strain, BFF23: CB1467 *him-5(e1490)* V*; zdIs13 (tph-1p::GFP)* IV, CB5362: *tra-2(ar221)* II*; xol-1(y9)* X, AM140: *rmIs132 (unc-54p::Q35-YFP)* I, NL2099: *rrf-3 (pk1426)*, EG7841: *oxTi302 [eft-3p::mCherry::tbb-2* 3'UTR + Cbr-*unc- 119(+)*].

## RNAi Treatment

We used a standard RNA interference (RNAi) feeding protocol, as previously described [19]. In each stage of the different experiments, worms were cultivated either on HT115 bacteria that transcribe a specific dsRNA (e.g., targeting *mCherry* or *dpy-11*) or on control HT115 bacteria that contain only an “empty vector” that does not lead to dsRNA transcription. The NGM plates contained IPTG for induction of dsRNA expression. The offspring which hatched on these plates were examined.

## Image Binarization, Cleaning & Segmentation

A mask image is produced by applying adaptive local thresholding [7], using the free parameters ‘neighborhood’ and ‘threshold’. The first determines the number of pixels around a given pixel that would be considered when deciding its value, while the second sets the relative intensity for thresholding the given pixel. The mask image is a matrix of equal size, as the original image, containing Ones where a suspected worm is present, and Zeros in the background. The filter’s parameters, ‘neighbors’ and ‘threshold’, may be adjusted manually, though several default recommended presets are offered, per the image’s resolution. After binarization, objects with areas smaller than a specified value will be deleted, including objects touching the edges of the image, which are unlikely to be whole worms. All the objects in an image are identified using MATLAB’s “regionprops” function, and are considered worms only if their area is within a specified relative range.

## Flagging Faulty Worms

Aberrant worms are flagged using two methods. First, the program locates outliers - worms with extremely large or small areas or lengths, or with non-continuous skeletons. Then, a C-NN network developed particularly for this task is used. This network has been trained on over 8000 mask images, half of valid worms and half with a variety of faulty features. New mask images are first padded and rescaled to fit the network, and then classified into “Worm” or “Non-Worm” categories, clearly marked for the users’ convenience.

## Morphological Measures

The calculation of the worms’ morphological features includes several steps. First, the worm is skeletonized to a single line using the “bwmorph” function, which is then cleaned and pruned from branches to obtain the worms’ straight skeleton. Afterwards, the worms’ edges are located using the “edge” function with the “Sobel” technique. A worm’s area is calculated using the “bwarea” function on the mask image, which is then multiplied by pixel height and pixel width. Its length is calculated by using the same function on the skeleton, which is then multiplied by pixel width. A pixel’s height and width is given as editable input in the program’s interface. Thickness is calculated by dividing the area by the length. Middle-width is computed using our own “cross_section” function, which locates the length of the worm’s cross section which is perpendicular to the pixel in the exact center of the skeleton. Lastly, the extraction of head and tail diameter ratios was adapted from WormGender [20], but with some modifications. The software calculates the mean intensity in the bright-field image around the two ends of the worm’s skeleton. It was previously shown [21] that ends with higher intensity are characteristic to the head of the worm. The first diameter of each end (D1) is the *longest* cross-section found 10 pixels from the worm’s end and up to 10% of the worm’s length. If the longest cross section is at 10% of its length, then the length of the cross-section at 2.5% of the worm’s length is taken as D1. The search for the second diameter of each end (D2) begins 20 pixels from D1’s location, and continues up to 20% of the worm’s length, until the *shortest* cross-section is found. Lastly, the diameter ratio for each end is calculated by dividing D1 by D2. We developed this algorithm to maximize the diameter ratio of the wider male tails [21], without biasing against hermaphrodites’ tails, to improve distinctiveness between sex phenotypes. Following these adjustments our software distinguishes the worms’ sex successfully in 95% of the cases tested (see in details bellow in the Results section).

## Fluorescent Analysis

We applied MATLAB’s “LocalMaximaFinder” to locate the local maximum intensities (Peaks) throughout the image, using the adjustable parameters – Neighborhood and Threshold. The ‘Neighborhood’ parameter specifies the size of a square surrounding an identified peak, within no other local maxima can be considered as peaks. A large Neighborhood allows only the brightest and most further apart peaks to be identified, while a small neighborhood allows many adjacent peaks to be identified separately. The ‘Threshold’ parameter enables control over the minimal intensity that can be considered as a peak, and is set by choosing the desired percentage from the image’s maximum intensity. The number of peaks identified, their mean intensity, and standard deviation are reported for each worm. In addition, Raw Integrated Density (RID) is calculated by summing the intensity values of all pixels within the worm’s area, and subtracting the mean background intensity multiplied by the number of pixels in the worm’s area. Lastly, Corrected Total Cell Fluorescence (CTCF) is calculated by subtracting the mean intensity within the worm from the mean intensity of its background, which is then multiplied by the worm’s area.

## Binary Classification

This allows the creation of a Support Vector Machine (SVM) model for binary classification, based on a labeled data set generated by the Feature Extractor. Users may choose a kernel method, whether to standardize their data, and the number of cross validations on the data, aimed at reducing model overfitting. The program splits the data to a training set and a test set, and performs optimization towards an appropriate SVM model. The resulting model may be later used to classify unlabeled data sets with similar features into the labels it was trained on. One may create one’s training data set by manually labelling worms, and then use the model created based on the data to classify unlabeled worms. Alternatively, one may utilize the data set to obtain prediction rates for various combinations of features, in order to assess the contribution of each feature towards accurate prediction (**Fig. 2**). We supply a trained model for the purpose of worm sex classification, but recommend customized labeling and training to create bespoke models that would better suit each lab’s specifications.

## Unsupervised Scoring

"t-Distributed Stochastic Neighbor Embedding" [4], is a technique for dimensionality reduction that is particularly well-suited for the visualization of high-dimensional datasets, and gives each data point a location on a two or three-dimensional map. The technique is a variation of Stochastic Neighbor Embedding that is easier to optimize, and produces better visualizations [22]. This method of unsupervised learning essentially enables scoring data within a single common dimension, giving each sample a continuous value. The data is pre-processed using “Principal Component Analysis” (PCA) [5], reducing the dimensionality. Later, dimensionality is reduced again via the t-SNE technique. If the data is indeed labeled, although labels are not used by t-SNE itself, they can be used to color the resulting plot. The algorithm’s final output is the low-dimensional data representation.

## Acknowledgements

We want to thank all the Rechavi lab members, and especially Moran Neuhof Dror Cohen, and Itamar lev for their help and for many discussions. Some strains were provided by the CGC, which is funded by NIH Office of Research Infrastructure Programs (P40 OD010440). A.Hakim and Y.Mor wish to thank the Sagol school of Neuroscience. Y.Mor is thankful for the TEVA NNE fellowship. A.Hakim is grateful to Dino Levi for his mentoring and support. O.R is thankful to the Adelis foundation, ERC grant #335624, and ISF grant #1339/17 for funding.

## Author contributions statement

A.Hakim and Yael Mor conceived, designed and developed the software. I.T helped with the design, experiments, and implementation of the program, Yishai Malkovich helped with imaging and analysis of the sexual phenotype assays. A.L performed the aggregation assays and contributed the data of his PolyQ experiments. All authors helped with the writing of the paper.

